# Epigenetic priming with valeric acid unlocks multi-stress resistance in plants by targeted histone deacetylase inhibition

**DOI:** 10.64898/2026.06.17.732912

**Authors:** Olaf Geyderowicz, Marta Gapińska, Helena Kossowska, Radosław Mazur, Patrycja Zembek, Roksana Iwanicka-Nowicka, Magdalena Krzymowska, Jarosław Poznański, Keqiang Wu, Łucja Kowalewska, Marta Koblowska

## Abstract

Progressive climate change is driving increasingly devastating crop losses through droughts, heatwaves and other adverse weather. Combined with shrinking arable land and spreading plant diseases, this makes it crucial to enable crops to grow efficiently in unfavorable, dynamic environments. Because stressors demand prompt, highly coordinated responses, plant stress adaptations rely heavily on epigenetic regulation. Histone deacetylases (HDAs), particularly class I, were recently shown to repress these responses. As constitutive defense is energetically costly, tools enabling temporal modulation of such mechanisms are highly sought after in crop biotechnology. This study evaluated whether valeric acid (VA), a five-carbon carboxylic acid, can inhibit HDA and activate plant defense responses.

Here we show that VA is a potent HDA inhibitor that confers resistance to multiple abiotic and biotic stresses in Arabidopsis, and further validate the abiotic component in maize and tomato. Despite its simple chemical structure, structural and transcriptomic evidence shows that VA acts by selectively inhibiting two major stress-repressing deacetylases, HDA19 and HDA6. By targeting these epigenetic switches, VA activates natural defense and acclimation responses. Time-course transcriptomic analyses further revealed that priming with VA induces transcriptional memory, which, together with VA-induced metabolic rewiring, enables rapid and efficient responses to future stressors. Most importantly, despite activating energetically demanding defenses, VA promotes vegetative growth and increases yield under normal conditions, thereby breaking the growth-defense trade-off.

These findings establish VA as the first epigenetic biostimulant of its kind, capable of improving plant performance under both abiotic and biotic stress while simultaneously increasing crop yield. As the epigenetic mechanisms underlying its action are evolutionarily conserved, VA priming emerges as a promising universal strategy to mitigate the climate-driven global crisis of crop losses.

## Introduction

Due to progressive climate changes, sustaining global food production and meeting the demands of growing population is, and will continue to be one of the most important challenges for plant scientists. Adverse weather conditions caused by rising global temperatures like drought and heatwaves cause increasingly more devastating crop yield loses (Razzaq et al. 2021). In addition, rising sea levels and the progressive desertification of arable lands more often expose plants to salt stress and it is presumed that they will cause global agricultural production to fall by 36.6% by the end of the 21st century (Hultgren et al. 2025). For this reason, it is crucial to find solutions that enable crops to be grown under unfavorable environmental conditions and render them resistant to periodic extreme weather events. Although most plants are sessile and unable to avoid environmental stress conditions, they have evolved sophisticated molecular mechanisms to adapt and survive under stress. As most of the responses are highly dynamic, often complex epigenetic mechanisms are behind orchestrating plant response to abiotic environmental stimuli (Chang et al. 2020).

Histone deacetylase (HDA/HDAC) are highly evolutionarily conserved enzymes involved in epigenetic regulation of gene expression in all eukaryotes (Alinsug, Yu, and Wu 2009). Their activity relies on the removal of acetyl groups on lysine residues of histone tails, increasing the affinity of nucleosomes to DNA leading to chromatin condensation and transcription silencing. In *Arabidopsis thaliana* there are 18 histone deacetylases (HDA) that can be divided into three families: 12 members of zinc dependent RPD3/HDA1 family, 4 members of plant-specific HD2 family and 2 members of Sir2 family, whose members use NAD as a cofactor. Based on phylogeny, the most diverse RPD3/HDA1 family can be further divided into three distinct classes based on their homology to yeast: class I (HDA6/7/9/19/10/17), class II (HDA5/8/14/15/18) and class III containing only one protein- HDA2 (Alinsug, Yu, and Wu 2009; Luo et al. 2017a; Minoru Ueda et al. 2017). Among the members of this family, class I enzymes are of greatest scientific interest due to their role in regulating responses to both biotic and abiotic stress (Hollender and Liu 2008; Luo et al. 2017b). Two class I deacetylases, HDA19 and HDA6, share overlapping roles in abiotic and biotic stress: both negatively regulate drought (Kim et al. 2017; M. Ueda et al. 2018; D. Xu, Leister, and Kleine 2024), repress SA-mediated defense genes (Jang et al. 2011; Y. Wang et al. 2017; Zheng et al. 2023), and act in salt stress response (L.-T. Chen et al. 2010; Minoru Ueda et al. 2017). HDA19 additionally represses heat response (M. Ueda et al. 2018) and JA response genes (C. Zhou et al. 2005). A third class I deacetylase, HDA9, is a crucial component of plant heat sensing (Niu et al. 2022), with knock-out mutants showing increased resistance to drought, salt stress, and *Pseudomonas syringae* pv. *tomato* DC3000 infection (Yang et al. 2020; Zheng et al. 2016). Taking into account highly resistant phenotypes of *hda19, hda6* and *hda9* mutants, class I HDA are promising but challenging targets in plant genetic engineering as well as other biotechnological approaches (M. Ueda et al. 2018; Minoru Ueda et al. 2017).

Although HDAs are of great importance in plant stress response modulation, most HDAs also play crucial roles in plant development, and their mutants often show various growth defects. For example, highly resistant *hda19* loss-of-function mutants show malformations of floral organs and near sterility (Gorham et al. 2018; Kotnik et al. 2025). Additionally, continuous maintenance of defense response is energetically demanding thus many mutants show stunted growth (Y. Wang et al. 2017). The solution to the abovementioned problems is the possibility of temporal control of HDA activity in plants which is possible with the use of histone deacetylase inhibitors (HDAi). Most of those compounds were developed primarily for the use in anticancer therapies in humans (Eckschlager et al. 2017), yet it has been shown that they are also functional in plants. Previous studies successfully reported that the use of Trichostatin A (TSA) or sodium butyrate (NaBT) inhibits HDA activity and enhances salt stress tolerance in Arabidopsis (Minoru Ueda et al. 2017). Additionally, rice plants treated with NaBT showed increased resistance to pathogenic fungus *Magnaporthe oryzae* in a recent study (Y. Xu et al. 2022). In many cases HDAis used in plant studies are of limited or lack selectivity which leads to complex, often unfavorable phenotypes. Thus, it has been lately postulated that there is a need for a selective HDAi that only affects stress responsive processes directed by HDA in plants (Sako, Nguyen, and Seki 2021). Studies on HDAi selectivity in human cell lines showed that small inhibitors of very simple chemical structure like sodium butyrate, regarded in many plant studies as non-selective (Davie 2003; Minoru Ueda et al. 2017), show selectivity towards class I HDA (Bantscheff et al. 2011; Schölz et al. 2015).

Valeric acid (VA) is a simple five-carbon aliphatic carboxylic acid that occurs abundantly in the natural environment. This compound is produced by certain plants, including *Valeriana officinalis* L., as well as by bacteria of the *Clostridia* and *Megaspharea* genera, which are part of the human intestinal microbiome (Markowiak-Kopeć and Śliżewska 2020). In agriculture, valeric acid can also be found in manure (Ni et al. 2012). Lately it has been shown that VA has the ability to inhibit HDAC3, human homolog of HDA19, in cancer cells (Han et al. 2022). Additionally, structurally similar compound, valproic acid (VPA, 2-propylvaleric acid), a commonly used drug in epilepsy treatment, shows unprecedented selectivity towards subset of class I HDA in humans (Bantscheff et al. 2011; Schölz et al. 2015). All the abovementioned data suggested that valeric acid would be a promising selective inhibitor of class I HDA that could potentially be used to epigenetically modulate plant stress response.

We therefore asked whether valeric acid, despite its simple structure, could function as a selective class-I HDA inhibitor. Unlike *hda* knock-out mutants, which exhibit enhanced stress resistance at the cost of impaired development and growth, a temporary chemical inhibitor could separate resistance from these drawbacks, offering a flexible alternative to genetic engineering for future crop protection. Our results show that VA acts as a selective class-I HDA inhibitor that provides resistance to various abiotic and biotic stresses in Arabidopsis, with similar phenotype observed in maize and tomato. Analysis of the resistant phenotype reveals that epigenetic priming with VA changes the plant’s energy metabolism and induces a stress memory that allows for better responses to subsequent stressors. Furthermore, VA priming breaks the growth-defense trade-off and increases yield under normal conditions.

This study focuses on three questions: how - what is the molecular mechanism of VA action; what - what phenotype does VA induce; and why - what cellular and systemic changes underlie resistance.

## Materials and Methods

### Plant materials and growth conditions

*Arabidopsis thaliana* accession Columbia (Col) was used if not indicated otherwise. The seeds were surface sterilized and sown on half-strength Murashinge and Skoog (MS)(Murashige and Skoog 1962) medium with Gamborg vitamins (Gamborg, Miller, and Ojima 1968) on Petri plates. After 48h stratification at 4°C in darkness, the plates were transferred to a growth chamber with 150 µmol × m^-2^ × s^-1^ 16h long photoperiod and 22°C. The plants were grown under those conditions for the desired amount of time before being used in in vitro stress experiments or being transferred to soil.

For the tests on maize, *Zea mays* L. grains were sown on moist presterilized sand and incubated for 5 days at 37°C in darkness. After this period, the seedlings were transferred either to soil or hydroponics installation for further growth and stress treatments.

In the case of tomato (*Solanum lycopersicum*) the seeds were germinated in pots with soil (9×9 cm, one per pot) and after 4 weeks the plants were directly used in the experiments.

### Heat stress treatment

In vitro *test on Arabidopsis thaliana-* 14-day-old seedlings were treated with sodium valerate by foliar or in-medium application. For foliar application, plates were sprayed with 1/10/100 mM VA or water (control); for in-medium application, plants were moved to half-strength MS with 100 µM/1 mM/10 mM VA or no inhibitor. After 3 days, plates were heat-shocked at 43.5°C (CLN53, POL-EKO) for 4 h according to (M. Ueda et al. 2018). Plants then regenerated at 22°C under long days for 7 days, after which plates were photographed and survival rate (% alive) determined. The schematic workflow of heat shock treatment is shown in Fig. S1b.

*Test on hydroponically grown maize*- germinated 5-day-old *Z. mays* seedlings were transferred to pots (16 per pot) with constantly aerated Knop’s solution and kept at 26°C under long-day greenhouse conditions (∼250 µmol × m^-2^ × s^-1^ light). Upon reaching the V3 stage, sodium valerate was added to the medium to the final concentration of 1 or 10 mM and the plants were grown for additional 3 days. After this time plants were transferred to fresh pots with identical media and heat-shocked at 50°C for 4 h, then returned to the greenhouse. After 7 days, leaves were excised at the base, photographed, and their necrotic area measured.

### Freezing stress treatment

14-day-old Arabidopsis plants were treated with NaVA in medium for 3 days as described above. The freezing test was performed in accordance with (Xin and Browse 1998) (Fig. S1d). After two days the plants were photographed, and the survival rate was determined.

### Salt stress treatment

7-day-old Arabidopsis seedlings were treated with sodium valerate in medium for 3 days, then transferred onto matching MS media (with or without VA) containing 200 mM NaCl. After 7 days, plates were photographed and survival rate determined.

### Drought stress treatment

*Arabidopsis thaliana*- 14-day-old plants were transferred to pots (9×9 cm, five per pot) with soil pre-wetted with VA (100 µM / 1 mM / 10 mM) or water. Plants were grown in greenhouse conditions (150 µmol × m^-2^ × s^-1^ 16h long photoperiod; 22°C; 60% relative humidity) for 10 days. After this period the soil in pots was dried using paper towels and the 16-day-long period of water deprivation began. At the end of this period the pots were photographed, and plants were rehydrated. After 2 days upon rewatering the pots were photographed again, and survival rate was determined.

*Zea mays*- germinated 5-day-old maize seedlings were transferred to pots with soil (14×14 cm, five plants per pot) and grown under greenhouse conditions (250 µmol × m^-2^ × s^-1^ 16h long photoperiod; 26°C; 60% relative humidity) for 14 days. After this time the remaining water was removed using paper towels and aqueous solution of sodium valerate (1 mM/ 10 mM) or pure water was added to each pot. Following the application of VA the plants were left for 16 days without additional watering for drought treatment. At the end of drought period plants were rewatered and after 5 days they were photographed and survival rate was determined.

*Solanum lycopersicum*- after 4 weeks of growth, remaining water in soil was removed using paper towels and aqueous solution of sodium valerate (0.1 / 1 / 10 mM) or pure water was added to each pot. The plants were photographed after 7 days without watering.

### Transpiration rate measurements

The protocol for water loss determination was adapted from (Cai et al. 2024). Briefly, 14-day-old Arabidopsis seedlings grown 3 more days on half-strength MS with various sodium valerate concentrations were used. Sixteen plantlets per condition were placed on Whatman paper and their pooled mass was measured on a precision scale at intervals over 90 min at 22°C and 40% relative humidity; fresh-weight loss rate was taken as transpiration rate.

### Stomatal Aperture Index measurements

The physiological measurements of stomatal aperture were performed according to (Eisele et al. 2016). For analysis, 14-day-old Arabidopsis seedlings were transferred onto half-strength MS media without or with various concentrations of sodium valerate for 3 days. After this time, the third youngest leaves from 5 plants from each plate were detached and floated on stomatal opening buffer (10 mM MES, 10 mM KCl, 0.1 mM CaCl_2_, pH=6.15). After 2 h of incubation under 150 µmol × m^-2^ × s^-1^ light at 22°C, abscisic acid (ABA)(Sigma Aldrich) was added to the buffer to a final concentration of 10 µM and leaves were incubated for additional 60 or 120 minutes. In each timepoint (2/3/4 h) the abaxial epidermis was peeled off and analyzed under light microscope (Axioscope 5, Zeiss). For image analysis and determination of Stomatal Aperture Index (SAI = aperture width / aperture length) of single stomata Fiji (Schindelin et al. 2012) was used. For each condition and each timepoint, roughly 100 individual stomata were analyzed.

### Infection with Pseudomonas syringae pv. tomato DC3000

For biotic stress experiments, 14-day-old Arabidopsis seedlings were transferred onto half-strength MS with or without 1 mM sodium valerate for 3 days, then dip-inoculated with *Pseudomonas syringae* pv. *tomato* DC3000 (culture density 10^8^ or 10^7^ CFU/ml). After infection, plants were placed back on the MS media and transferred to the growth chamber. To measure bacterial growth, samples were collected at 0, 2, and 4 days post-infection (dpi). Three plants per plate were pooled at each time point. Shoots were separated from roots, surface-sterilized in 70% ethanol for 30 seconds, rinsed with sterile water, blotted dry, and weighed. Tissue was then homogenized in 300 µl of 10 mM MgCl₂.

Serial dilutions were plated on LB agar supplemented with 50 µg/ml rifampicin for bacterial enumeration. After two days of incubation, colonies were counted, and CFU per mg of shoot tissue was calculated. The experiment was conducted in four replicates.

### RNA isolation

For transcriptomics under normal conditions, four whole plants per plate were pooled and frozen in liquid nitrogen 24 and 72 h after 14-day-old seedlings were transferred onto half-strength MS with 1 mM VA. For the studies under heat stress (HS) conditions, plates with plants growing for 72 h on 1 mM VA supplemented medium were transferred to an incubator set to 43.5°C for 4 hours. Incorporating the same pooling model, plants were frozen immediately after the end of HS treatment and 24 hours later. The whole experiment was performed in four replicates. 100 mg of each frozen sample was used as starting material for RNA isolation using E.Z.N.A.® Plant RNA Kit (Omega Bio-Tek).

### Microarray analysis

For microarrays, the highest-quality RNA from three biological replicates was processed independently. Reverse transcription, cRNA synthesis, second cycle ss-cDNA synthesis, fragmentation and labeling were performed using GeneChip™ WT PLUS Reagent Kit (Applied Biosystems). After labeling, the samples were hybridized to GeneChip™ Arabidopsis Gene 1.1 ST Array Strips (Applied Biosystems), scanned using GeneAtlas Scanner (Affymetrix). For data analysis the files were uploaded to Transcriptome Analysis Console (TAC) Software (ThermoFisher) where background correction was carried out and probe signals were calculated using RMA algorithm (Irizarry 2003). After data normalization and log_2_ transformation, Principal Component Analysis was carried out to identify potential outliers. In order to obtain lists of Differentially Expressed Genes (DEGs) eBayes ANOVA (Ritchie et al. 2015) was performed with the following cutoffs: fold change (FC) ≥ 2; p-value < 0.05.

The complete datasets of the microarray experiment are available in the NCBI Gene ExpressionOmnibus (GEO) database repository with accession number GSE316766.

### Transcriptomic data analysis

Functional enrichment analysis, based on GeneOntology (GO) and Kyoto Encyclopedia of Genes and Genomes (KEGG) databases, for DEGs in each timepoint was conducted using STRING v.12 server (Szklarczyk et al. 2023). In order to identify functional hubs in the network, MCL clustering (inflection parameter = 3) was applied and functional enrichment for each cluster has also been separately performed. In order to identify transcription factors in DEGs lists, PlantTFDB v.5.0 was used (Jin et al. 2014). Complimentary analyses were also performed using Metascape (Y. Zhou et al. 2019). In all cases up and down-regulated genes were analyzed separately.

### Real Time qPCR assay

To validate the transcriptomic data, RT-qPCR was performed on the microarray RNA samples. After reverse transcription (RevertAid First Strand cDNA Synthesis Kit, ThermoFisher), RT-qPCR was run on a LightCycler® 96 (Roche) with SYBR™ Select Master Mix (Applied Biosystems). Primer sequences are listed in Supplementary Table 1.

### Quantification of histone acetylation levels by Western Blotting

For histone isolation, 200-600 mg frozen Arabidopsis seedlings were used and procedures were performed as previously described (Przewloka and Jerzmanowski 1997). Proteins were separated by SDS-PAGE electrophoresis. The histone modifications were detected by immunoblotting using the following primary antibodies: anti-H3 (Sigma Aldrich, 9289), anti-H3ac (Upstate, 06-599), anti-H3K9ac (Sigma Aldrich, H9286), anti-H4K5ac (Diagenode, C15410025) and secondary HRP-conjugated anti-rabbit IgG (A9169, Sigma-Aldrich). The detection was performed using the AgriseraECL SuperBright (Agrisera) according to manufacturer’s protocol.

### MAL assay for HDA activity

Enzymatic tests were performed using a synthetic fluorescent histone deacetylase substrate, MAL (Boc(Ac)Lys-AMC) (Sigma Aldrich) (Heltweg et al. 2003). The tests were performed on protein extracts from *Arabidopsis thaliana* T87 suspension cultures. Two grams of frozen 7-day-old cells were ground in liquid nitrogen and suspended in isolation buffer (20 mM Tris-HCl pH 8, 10 mM NaHCO_3_, 10 mM MgCl_2_, 1 mM EDTA, 5 mM DTT, 1×cOmplete® Protease Inhibitor Cocktail (Roche)) in a 1:3 ratio. The suspension was filtered through a double layer of Miracloth membrane (Milipore) and dry (NH_4_)_2_SO_4_ was added to 90% saturation to precipitate the proteins. After 30 minutes incubation on a laboratory rotator at 4°C, the suspension was centrifuged at 18,500×g at 4°C for 20 minutes. The supernatant was decanted and the pellet was resuspended in E1 buffer (1.4 mM NaH_2_PO_4_, 18.6 mM Na_2_HPO_4_, pH 7.9, 0.25 mM EDTA, 10 mM NaCl, 10% (v/v) glycerol, 10 mM 2-mercaptoethanol) to obtain a total protein concentration of 10 mg/ml. The extract prepared in this way was used as the starting preparation for the assessment of histone deacetylase enzymatic activity. The assay was performed according to the original publication (Heltweg et al. 2003). The final concentrations of sodium valerate or NaBT in the reaction mixtures ranged from 1.6 µM to 83 mM. As a reference inhibitor, Trichostatin A (TSA) (Sigma Aldrich), a known non-specific inhibitor of histone deacetylases, was used in concentrations ranging from 3.2 nM to 50 µM. Fluorescence (330/390 nm) measurements were conducted using a CLARIOstar Plus reader (BMG Labtech).

### Starch quantification

Starch content was measured as described in (Smith and Zeeman 2006). 100-200 mg of frozen Arabidopsis seedlings were boiled in 80% ethanol at 100°C for 5 min, centrifuged at 5,000×g for 5 min, and the supernatant discarded; this wash was repeated twice. The pellets were then homogenized in distilled water and heated to 100°C for 10 minutes. One volume of 200 mM CH_3_COONa (pH=5.5), 7U of amyloglucosidase (Sigma Aldrich) and 3U of amylase (Sigma Aldrich) were added to the cooled samples, and the reaction mixtures were incubated at 37°C overnight. After incubation the glucose concentration in each sample was determined using assay with hexokinase (Sigma Aldrich) and glucose-6-phosphate dehydrogenase (Sigma-Aldrich). Produced NADPH was quantitated using a Cary50Bio spectrophotometer.

### Metabolic profiling

One hundred to 200 mg frozen Arabidopsis seedlings were ground in liquid nitrogen and the powder resuspended in 1.5 ml of chloroform : methanol : water (2 : 1 : 0.8) and thoroughly mixed. The suspension was centrifuged at 10,000×g for 5 minutes. 1 ml of the supernatant was then transferred to a new microcentrifuge tube, 500 µl water was added and the mixture was vigorously mixed to facilitate extraction. After phase separation the upper phase containing polar metabolites was taken for further analysis using an ion-exchange chromatography system (Dionex) coupled with Waters ZQ mass spectrometer as described in (Owczarek et al. 2020) with slight modifications.

### Photosynthetic Activity Measurements

Chlorophyll fluorescence was measured using the MAXI version of the Imaging-PAM chlorophyll fluorescence system (Heinz Walz, Germany) as described in (Podgórska et al. 2020). Petri plates with Arabidopsis plantlets were dark-adapted for 30 minutes before the experiment. After Fv/Fm (maximal efficiency of PSII photochemistry in the dark-adapted state) measurement, the blue actinic light of 200 µmol × m^-2^ × s^-1^ was applied for 5 minutes, and saturation pulses (SP) for Fm’ and F_0_’ determination were applied each 20 seconds., From the last SP, the effective photochemical yield of PSII (Y(II)), regulated non-photochemical yield of PSII (Y(NPQ)), and non-regulated non-photochemical yield of PSII(Y(NO)) were calculated using ImagingWinGigE software using the equations from (Kramer et al. 2004).

### Ultrastructural analysis using TEM

Plant specimens for ultrastructural analysis were prepared as previously described (Wójtowicz et al. 2025). The ultrathin sections (70 nm) were examined with a JEM 1400 electron microscope (Jeol, Japan). For data analysis, GRANA (Bukat et al. 2025) software was used to asses plastid ultrastructure. For other measurements and analyses Fiji software (Schindelin et al. 2012) was applied.

### Heterologous expression and purification of recombinant HDA19-GST

In order to obtain expression construct, full-length CDS of HDA19 (AT4G38130), amplified from *Arabidopsis thaliana* Col-0 total cDNA, was cloned into pGEX4-T1 vector. The resulting construct was transformed into *E. coli* Rosetta 2(DE3) (Novagen). For protein expression the cells were grown in liquid LB medium supplemented with ampicillin and chloramphenicol under constant agitation at 37°C. Upon reaching OD_600_=0.6, IPTG was added to the final concentration of 0.1 mM and the cultures were left overnight at 30°C. HDA19-GST was then purified using Glutathione Sepharose™ 4B (Cytiva) according to manufacturer’s protocol.

### Low Volume Differential Scanning Fluorimetry (nanoDSF) measurements

For nanoDSF, the GST tag was cleaved with thrombin and HDA19 purified by inverse affinity chromatography. Protein was diluted to 0.2 mg/ml in Z1 buffer (150 mM HEPES pH=7.4, 250 mM NaCl, 5% glycerol, 1 mM DTT) and supplemented with 0.05% (v/v) Tween-20 to avoid aggregation. For analysis of HDA19-VA interactions sodium valerate was added to the samples to the final concentration of 50 mM. Samples prepared this way were loaded into nanoDSF grade Standard Capillaries (NanoTemper Technologies) and analyzed using the Prometheus NT.48 nanoDSF device (NanoTemper Technologies). The protein unfolding was monitored with laser excitation power of 80% and the temperature increment was set to 1°C/min (25-85°C). Data analysis was performed with MoltenProt v1.1 software (Kotov et al. 2021) on EMBL eSPC platform (Burastero et al. 2021) with equilibrium two-state model.

### Phylogenetic analysis of RPD3/HDA1 histone deacetylases

In order to analyze the phylogeny of histone deacetylase in *Arabidopsis thaliana* as well to find human homologs of distinct HDAs, UPGMA tree was constructed. Amino acid sequences of all 12 RPD3/HDA1 family histone deacetylases from Arabidopsis and 11 from *Homo sapiens* were obtained from UniProt database. After Multiple Sequence Alignment (MSA) using ClustalX2 (Larkin et al. 2007) the phylogenetic tree was constructed with UPGMA algorithm using PAUP4.0a (Swofford 2002).

### Inverse molecular docking

For inverse molecular docking using AutoDock Vina 1.2.0 (Eberhardt et al. 2021) structures of all 12 Arabidopsis RPD3/HDA1 histone deacetylases were obtained using AlphaFold3 (Abramson et al. 2024). 3D models of sodium valerate (3669864) and Trichostatin A (444732) were obtained from PubChem database (https://pubchem.ncbi.nlm.nih.gov/). Both for ligand and for receptor preparation the Meeko python package (Santos-Martins et al. 2025) was used. The performed docking was targeted on the active sites of the enzymes (grid box of 20×25×25 points, spacing 0.375 Å) and the exhaustiveness parameter was set to 32.

### hda mutants transcriptomic analysis

Lists of Differentially Expressed Genes (DEGs) in *hda6* (*shi5*), *hda9-1* and *hda19* mutants were obtained from (Y. Wang et al. 2017) and (Shen et al. 2019) respectively. In both cases for RNAseq transcriptomic analyses, 10-day-old plants growing on half-strength MS medium were used and sequencing was performed using Illumina technology.

### Statistical analysis

Unless stated otherwise, experiments were run in triplicate. Multiple comparisons used one-way ANOVA with post-hoc Tukey HSD; different letters in figures and tables denote significant differences at p < 0.05. Pairwise comparisons used a two-sided t-test, with significance marked by asterisks per APA guidelines.

## Results

### VA specifically inhibits class I HDA

To assess VA functionality as a plant HDA inhibitor, biochemical deacetylase activity assays with the synthetic MAL substrate were performed on protein extracts from *Arabidopsis thaliana* T87 suspension cultures.

(Fig. 1a-c). The pan-HDA inhibitor trichostatin A (TSA) confirmed strong non-specific activity, with an IC_50_ of 87.44 nM and full inhibition at 10 µM. VA inhibited HDA activity only above 100 µM and failed to reach complete inhibition even at 100 mM. The same pattern held for sodium butyrate (NaBT), a structurally similar HDA inhibitor widely regarded as non-specific (Minoru Ueda et al. 2017). This indicates that, despite their simple structure, both VA and NaBT are class-selective, inhibiting only part of cellular HDA activity at working concentrations (0.1-10 mM). VA nonetheless showed a lower IC_50_ value (33.41 mM) than NaBT (86.34 mM).

**Fig. 1.**
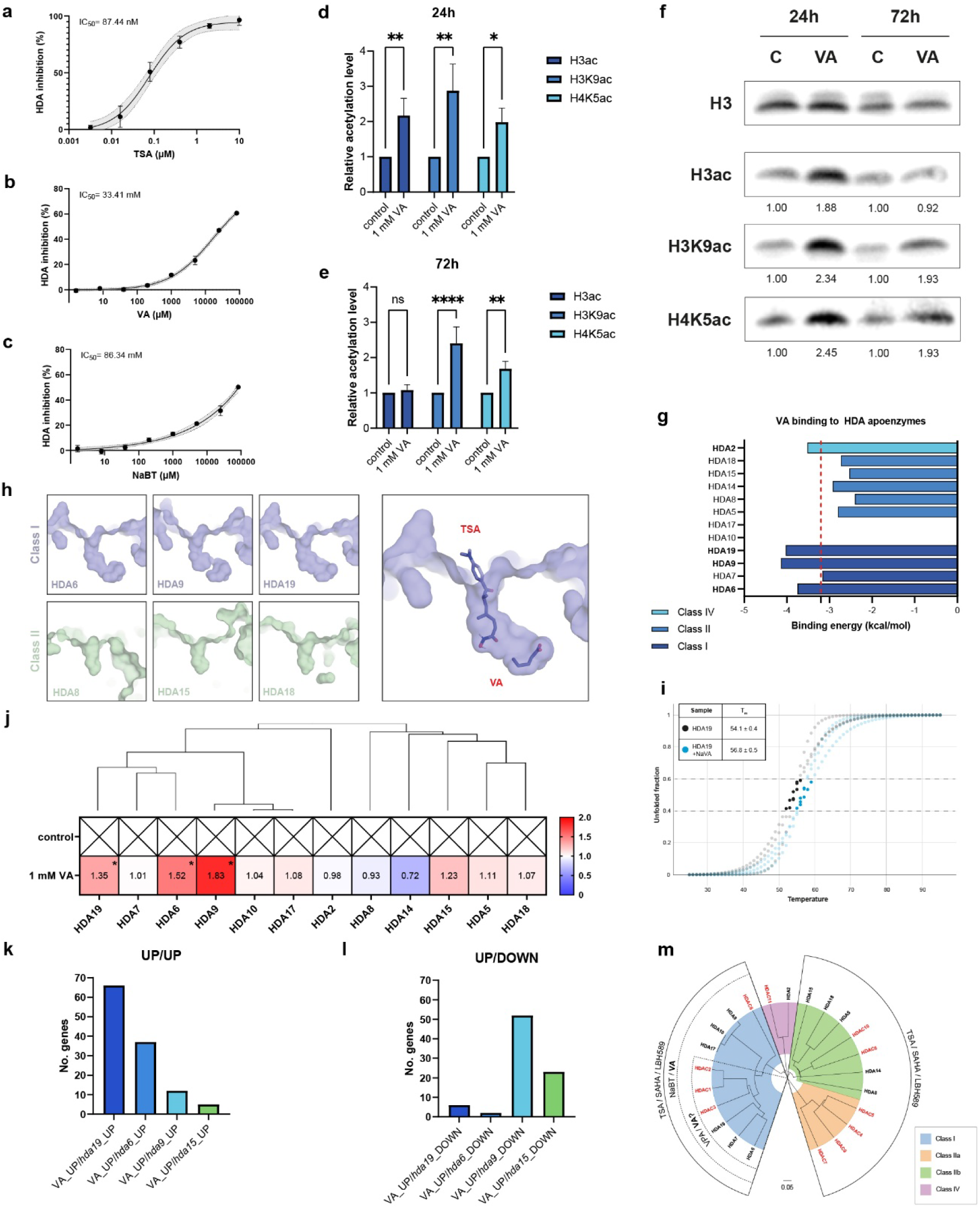
VA specifically inhibits class I HDA- HDA6 and HDA19. **a**, **b**, **c**, HDA dose-response inhibition curves for pan-HDA inhibitor trichostatin A (TSA), VA and sodium butyrate (NaBT) obtained on whole-cell protein extracts from T87 cell culture. **d**, **e**, **f**, relative levels of H3ac, H3K9ac and H4K5ac determined by western blot analysis after 24 h and 72 h of 14-day-old seedlings exposure to VA quantified by densitometric analysis. **g**, results of inverse docking using AutoDock Vina of VA to apoenzymes off all RPD3/HDA1 family histone deacetylases; red line denotes mean binding energy. **h**, structural comparison of the active site cavities of representative class I and class II histone deacetylases from *Arabidopsis thaliana* (left) and best score docking poses of TSA and VA to the HDA19 active site (right). **i**, nanoDSF unfolded protein fraction curves showing HDA19 thermal stabilization in the presence of 50 mM VA, indicating binding. **j**, phylogenetically clustered Arabidopsis RPD3/HDA1 deacetylases with their relative expression levels after 24 h VA treatment overlaid. **k**, number of common upregulated genes in VA-treated plants and *hda19, hda6, hda9* and *hda15* knock-out mutants. **l**, number of common genes that were upregulated in VA-treated plants but downregulated in *hda19, hda6, hda9* and *hda15* knock-out mutants. **m**, phylogenetic tree of RPD3/HDA1 deacetylases from Arabidopsis (black) and human (red) with the HDA inhibitor’s specificity (according to literature and this study) overlaid. Bars represent mean ±SD, with error bars showing variability among biological replicates. Data in (**a**, **b, c, d, e, f, i** and **j**) *n* = 3. All statistical tests were carried out using a one-way ANOVA with post-hoc Tukey test with asterisk (APA guidelines) indicating significant differences (*P* < 0.05). Source data are provided as a Source Data file.

To link HDA inhibition to histone acetylation, western blotting on histones isolated from VA-primed plants was performed (Fig. 1f). After 24 h of priming, VA doubled pan-acetylation of histone H3 tails (Fig. 1d). By 72 h this H3ac rise was no longer detectable, yet acetylation at specific stress-related lysines (Hu et al. 2019) remained elevated (Fig. 1e), indicating two distinct effector phases: an early response (24 h) and a late response (72 h).

Inverse molecular docking further resolved VA specificity: VA preferentially binds class I HDA (Fig. 1g). The weak or absent interaction with HDA7/10/17 likely reflects their being enzymatically inactive (Saharan et al. 2024). Structural analysis of Arabidopsis HDA showed that class I enzymes have an extended active-site pocket with an additional terminal cavity (Fig. 1h), to which VA preferentially binds, explaining its class I specificity. VA binding to the representative class I enzyme HDA19 was also confirmed in vitro by nanoDSF (Fig. 1i).

Transcriptomic analysis (Fig. 1j) showed that VA elevates expression of all enzymatically active class I HDAs (HDA6, HDA9 and HDA19). To test whether this reflects compensation for their inhibition, our data were compared with those of *hda6, hda9, hda19* and *hda15* (negative control) knock-out mutants. VA-induced changes align with those of *hda6* and *hda19* mutants, sharing many upregulated genes (Fig. 1k). Unexpectedly, more common DEGs were found between downregulated than upregulated genes in *hda9* and VA-upregulated ones (Fig. 1l), suggesting that elevated HDA9 expression is an indirect effect rather than direct compensation for its inhibition. VA is therefore specific not only to class I HDA but, within this class, to HDA6 and HDA19. As shown in Fig. S1a, over half of the VA-induced transcription factor changes are attributable to HDA6 and HDA19 inhibition. Figure 1m summarizes the revised specificity of common HDA inhibitors and VA on the deacetylase phylogenetic tree (Bantscheff et al. 2011; Schölz et al. 2015).

### VA unlocks broad resilience to abiotic stress

To test whether HDA6/HDA19 inhibition by VA enhances survival under abiotic stress, multiple in vitro and soil assays were performed.

For heat stress, the heat-shock protocol of (Ueda et al. 2018) was used. Three days of priming on 100 µM VA raised survival after a 4 h heat shock from ∼1% in controls to ∼23% (Fig. S1c), and 1 mM VA increased it further to ∼80%, while 10 mM in medium was toxic. Foliar application required 10-fold higher concentrations for comparable protection, consistent with poorer uptake through leaves than roots, plants sprayed with 10 mM VA reached the highest survival of all experiments (>95%) (Fig. 2a-b). The same heat protection was reproduced in hydroponically grown maize (Fig. 2d).

**Fig. 2.**
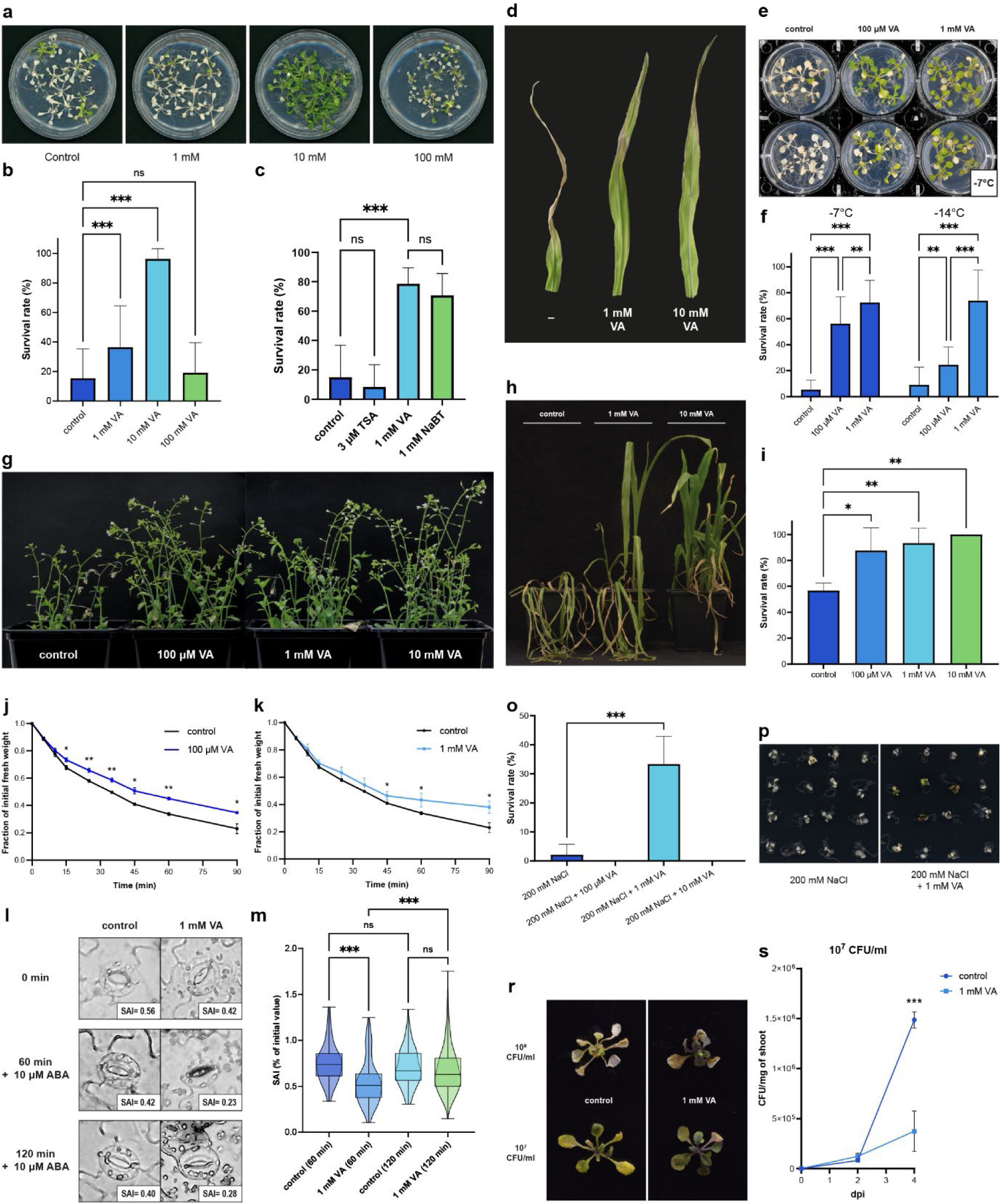
VA unlocks broad resilience to abiotic and biotic stress. **a**, appearance of Arabidopsis plants sprayed with aqueous solutions of different VA concentrations 7 days after heat shock (4 h, 43.5°C). **b**, survival rate of plants from different variants 7 days after heat shock. **c**, survival rate of plants treated with 3 µM TSA, 1 mM NaBT or 1 mM VA 7 days after heat shock. **d**, appearance of leaves of maize grown hydroponically with different VA concentrations 7 days after heat shock (4h, 50°C). **e**, appearance of Arabidopsis plants grown on MS media with various concentrations of VA 2 days after freezing to −7°C. **f**, survival rate of plants from different variants 2 days after freezing to −7°C or −14°C. **g**, **h**, appearance of Arabidopsis and maize plants watered with VA aqueous solutions after 14 days of drought followed by 7 days of rewatering. **i**, survival rate of Arabidopsis plants from different variants 7 days after rewatering preceded by drought period. **j**, **k**, transpiration rate of plants treated with 100 µM or 1 mM VA as compared to control. **l**, exemplary images of stomata from VA-treated and control plants upon ABA administration. **m**, stomata movement dynamics in control and VA-treated Arabidopsis plants. **o**, survival rate of Arabidopsis plants from different variants after 7 days of growth on media containing 200 mM NaCl. **p**, appearance of seedlings from different variants after 7 days of growth on high-NaCl media. **r**, Appearance of Arabidopsis plants grown on medium supplemented with 1 mM VA or devoid of it 4 days after dip inoculation with *Pseudomonas syringae pv.* tomato DC300 of given CFU/ml. **s**, number of bacterial cells over time in plants inoculated using 10^7^ CFU/ml *Pst* inoculum. Bars represent mean ±SD, with error bars showing variability among biological replicates. Data in (**b, c, o**) (*n* = 3, 16 plants per each replicate), in (**f**) (*n* = 6, 5 plants per each replicate), in (**i, j, k**) (*n* = 3, 5 plants per each replicate), in (**m**) (60 min: control *n*= 72, 1 mM VA *n* = 100; 120 min: control *n*= 158, 1 mM VA *n* = 225) in (**s**) (*n* = 3, 4 plants per replicate). All statistical tests were carried out using a one-way ANOVA with post-hoc Tukey test with asterisk (APA guidelines) indicating significant differences (*P* < 0.05). Source data are provided as a Source Data file.

It was also tested whether TSA and NaBT increase Arabidopsis survival after heat shock (Fig. 2c). NaBT increased heat resistance similarly to VA, albeit slightly weaker, whereas the pan-HDA inhibitor TSA was no different from control, showing that only selective class I HDA inhibition confers heat resistance in Arabidopsis.

Cold-stress experiments assessed resistance to sub-zero temperatures after VA priming (Fig. 2e). Priming with 100 µM VA markedly increased survival, from under 10% in controls to roughly half at −7 °C and a quarter at −14 °C (Fig. 2f). At 1 mM VA, survival rose to ∼70% regardless of freezing temperature, showing that VA confers resistance to both heat and cold.

Drought, another economically important stress, was tested in soil. Arabidopsis plants primed with aqueous VA before a 14-day drought survived markedly better than water-treated controls (Fig. 2i), and 10 mM VA gave full protection with complete survival and recovery. This tracked with reduced transpiration in VA-treated plants (Fig. 2j,k) and with narrower stomatal apertures under 1 mM VA than in controls (Fig. 2l). These stomata also responded faster to the abscisic acid closure signal, limiting water loss (Fig. 2m). The same drought-resistant phenotype was reproduced in maize (Fig. 2h) and tomato (Fig. S1e).

VA also conferred salt-stress resistance. Seedlings primed with 1 mM VA grew in the presence of 200 mM NaCl, whereas untreated plants did not survive (Fig. 2o, p). As in other in vitro tests, 10 mM VA was toxic, an effect never seen at this concentration in soil.

### VA confers resistance to biotic stress

To test whether VA also affects biotic stress responses, Arabidopsis infections with *Pseudomonas syringae* pv. *tomato* DC3000 were performed. As *P. syringae* is a hemibiotroph, it models both biotrophic and necrotrophic infection. At both inoculum densities (10^7^ and 10^8^ CFU/ml), VA-treated and untreated plants differed in disease severity (Fig. 2r). Bacterial counts were similar at 0 and 2 days post-infection (dpi), but by 4 dpi plants on 1 mM VA carried markedly fewer bacteria per mg shoot at the 10^7^ CFU/ml inoculum (Fig. 2s). VA therefore acts not by blocking early pathogen entry but by limiting colonization and disease progression in the tissues. This reflects primed plant defenses rather than direct antibacterial action, as VA at the concentrations used did not affect growth of *Pseudomonas syringae* pv. *tomato* DC3000 (Fig. S1f).

### VA primes stress response on transcriptomic level

At 24 h, VA priming upregulated 273 and downregulated 158 genes (Fig. 3a). Among upregulated genes, Metascape network analysis revealed three functional hubs (Fig. 3c): a dominant energy-metabolism hub (fatty acid degradation, oxylipin, pyruvate and starch metabolism); a pathogen-defense hub centered on jasmonic acid (JA) and salicylic acid (SA) responses, with JA terms linked to wounding and bacterial response, SA terms to fungal response, and toxin and sulfur metabolism also present; and a smaller abiotic-stress hub built around responses to water deprivation and oxidative stress. Downregulated genes formed two hubs, one for cell wall biogenesis and modification and one for gibberellin response (Fig. 3d). These patterns were mirrored at the transcription factor level (Fig. 3b). The most abundant upregulated TFs were WRKYs, including JA-dependent WRKY51, SA-dependent WRKY40, and WRKY70 (acting in both pathways); these are key activators of biotic and abiotic stress responses (Phukan, Jeena, and Shukla 2016). Two ethylene-responsive factors were also found, ERF113 (abiotic) and ERF2 (biotic) (Fujimoto et al. 2000). Although this cluster’s highest GO term is biotic, it also contains two ZAT transcription factors, ZAT10 and ZAT12, crucial activators of abiotic stress response under salt, cold and light stress (Mittler et al. 2006). Among the VA-downregulated transcription factors there are multiple ones involved in cell wall biogenesis and its modifications.

**Fig. 3.**
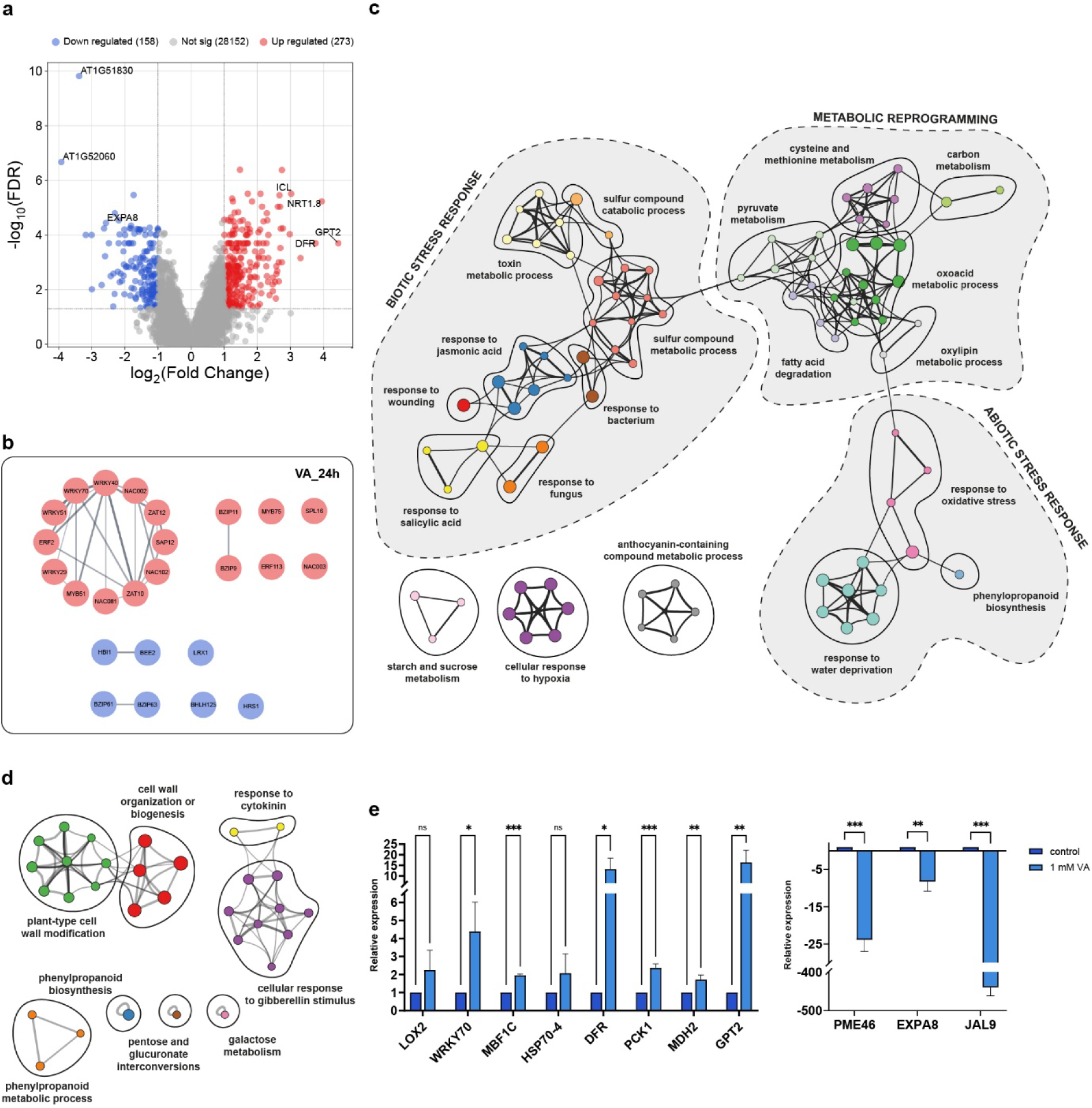
VA primes stress response on transcriptomic level. **a**, volcano plot showcasing global transcriptomic changes after 24 h of VA treatment. **b**, MCL clusters of transcription factors upregulated (red) and downregulated (blue) upon 24 h of VA treatment. **c**, GO functional analysis network of genes upregulated in 2-week-old Arabidopsis plants after 24 h of VA treatment. **d**, GO functional analysis network of genes downregulated after 24 h of VA treatment. **e**, RT-qPCR validation of chosen VA-upregulated (left) and VA-downregulated (right) genes identified in the microarray transcriptomic analysis. Bars represent mean ±SD, with error bars showing variability among biological replicates. Data in (**e**) (*n* = 3). All statistical tests were carried out using a one-way ANOVA with post-hoc Tukey test with asterisk (APA guidelines) indicating significant differences (*P* < 0.05). Source data are provided as a Source Data file.

Among all the clusters described above, representative genes were chosen and their expression levels in VA-treated and non-treated plants using RT-qPCR (Fig. 3e).

### VA-induced transcriptional memory under stress

To probe VA action under stress, heat shock, a well-established stress memory model, was used with sampling at 72 h after VA exposure (just before heat shock; VA_72h), immediately after heat shock (VA_4h_HS) and 24 h later (VA_28h_HS).

Venn diagrams of DEGs across timepoints (Fig. 4a-b) show that VA-primed plants respond dynamically, activating different gene subsets. Functional time-course analysis showed that genes upregulated after 3 days of VA resemble those at 24 h (Fig. 3c), with abiotic-stress defense, detoxification, and cell wall genes again downregulated (Fig. 4d). Four hours into heat stress, VA-treated plants showed elevated fatty acid biosynthesis (Fig. 4c), reflecting sustained JA biosynthesis and response, while SA response was quenched (Fig. 4d). Surprisingly, 24 h after heat shock, valerate-primed plants broadly activated the biotic stress response as SA-mediated systemic acquired resistance (SAR). This was accompanied by starch degradation and downregulation of heat-responsive genes, indicating faster stress recovery than in untreated plants (Fig. 4d).

**Fig. 4.**
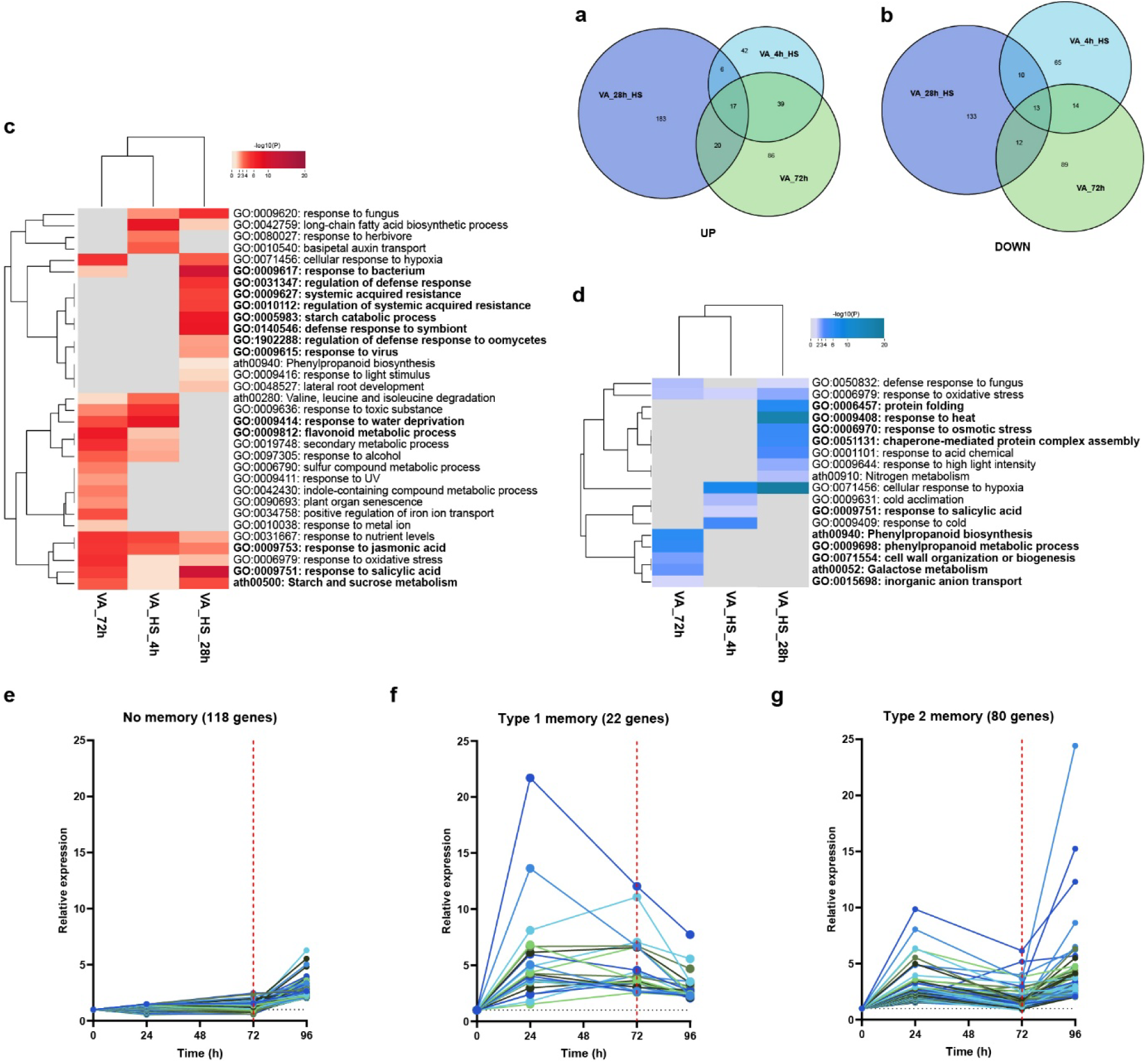
VA priming leaves transcriptional memory and helps plants to better respond during and after heat stress. **a**, **b**, Venn diagrams showing the number and overlap of upregulated and downregulated genes. **c**, **d**, Gene Enrichment heatmaps depicting functional changes among genes upregulated (red) and downregulated (blue) throughout the time course of heat stress experiment. **e**, **f**, **g**, line graphs depicting mean relative expression levels of genes upregulated 24 h after heat shock showing distinct transcriptional memory types. Source data are provided as a Source Data file.

This rapid biotic response, unrelated to heat stress, was attributable to epigenetic priming of those defense genes during the early response to VA. Almost half of the genes upregulated after heat shock had already been upregulated 24 h after VA priming. These 102 genes displayed both known types of transcriptional memory (Fig. 4e-g)(Oberkofler, Pratx, and Bäurle 2021). Twenty-two showed sustained induction (type-I memory) and were involved in metabolic reprogramming. The other 80 showed type-II memory (enhanced re-induction) and were the main drivers of the biotic response (Fig. S2). Transcripts upregulated only after heat shock (no memory) also contributed to the biotic response but likely play a secondary role. Thus, VA priming uses distinct memory types for distinct processes: type-I for long-term metabolic reprogramming and type-II for rapid responses to future stress.

Together, these findings show that upregulation of stress-response genes during the first 24 h of VA priming leaves transcriptional memory, potentially via locus-specific hyperacetylation (Harris, Amtmann, and Ton 2023). With accumulated energy reserves, this memory enables rapid, robust defense upon contact with a future stressor.

### VA-induced multi-stress resistance relies on metabolism reprogramming

Given the metabolism-oriented transcriptomic results, the VA-primed phenotype was analyzed by LC-MS metabolic profiling and biochemical assays.

Profiling showed that all measured TCA-cycle intermediates rose in VA-primed plants, with citrate/isocitrate, succinate and malate increasing most. This selective overaccumulation of only a few intermediates argues against general Krebs-cycle upregulation. Integrating transcriptomic data (Fig. 5a) revealed elevated expression of the glyoxylate-cycle markers isocitrate lyase (ICL), cytoplasmic malate dehydrogenase (cMDH2) and citrate synthase (CSY2) (Eastmond and Graham 2001) indicating activation of this alternative catabolic cycle. The glyoxylate cycle is normally active only in germinating seeds, to use acetyl-CoA from storage fatty acid degradation, with little being known on its function in mature plants. Oxaloacetate levels stayed largely unchanged, due to overexpression of phosphoenolpyruvate carboxykinase 1 (PCK1) promoting gluconeogenesis, process also facilitated by the upregulated pyruvate-phosphate dikinase 1 (PPDK). Carbohydrate accumulation was also evident transcriptomically: the most strongly induced transcript was the chloroplastic glucose-6-phosphate transporter (GPT2), which imports glucose-6P and other sugars into the stroma for starch synthesis. VA-treated plants began accumulating starch within 24 h and, after 3 days, contained several-fold more starch than controls (Fig. 5c). TEM images (Fig. 5b) show the corresponding chloroplast differences.

**Fig. 5.**
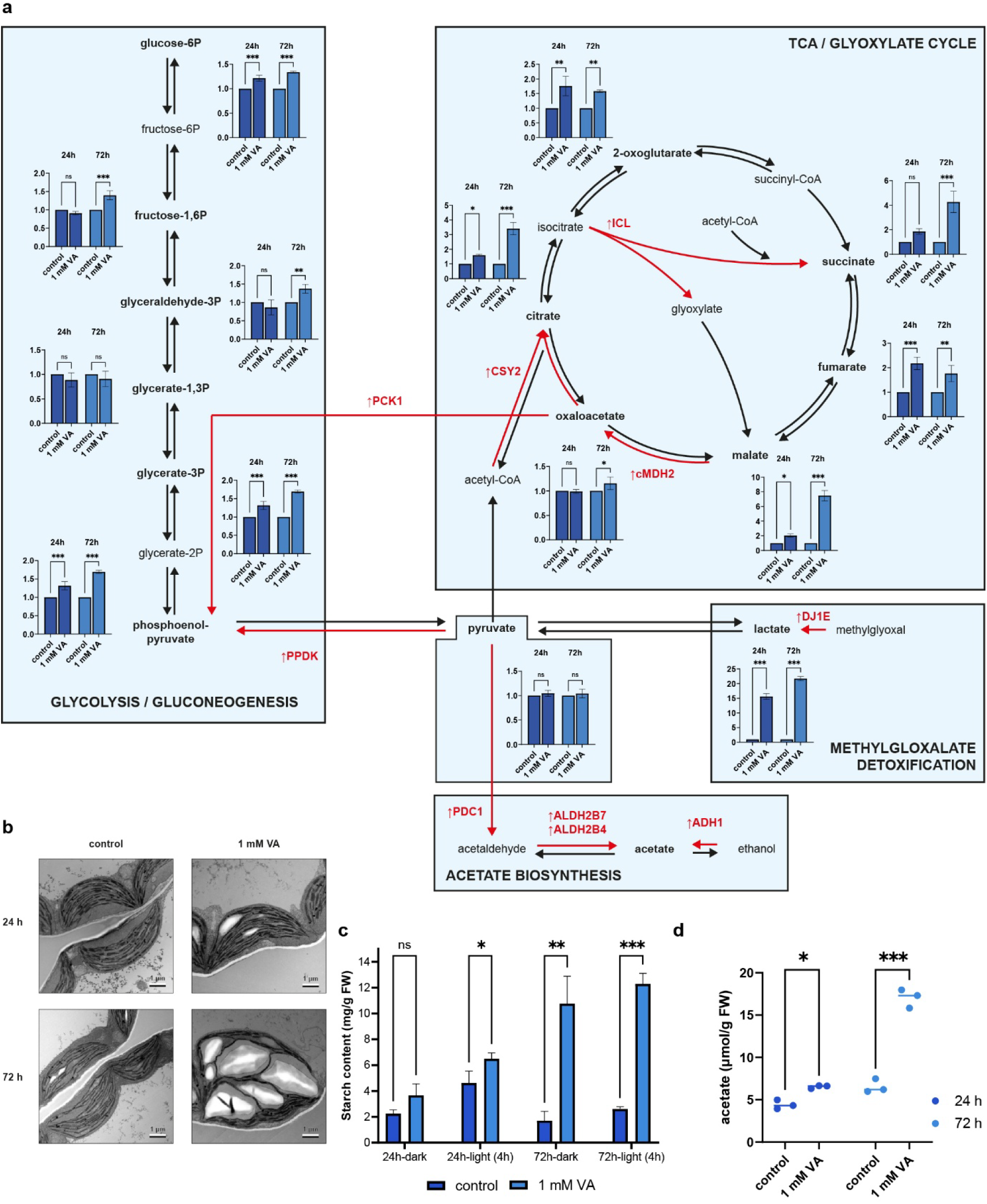
VA-induced resistant phenotype relies on major metabolism reprogramming. **a**, relative levels of glycolysis/gluconeogenesis, TCA/glyoxylate cycle, lactate and acetate biosynthesis metabolites quantified using LC-MS in VA-treated plants. In red are the names of enzymes which expression is induced by valerate treatment. **b**, exemplary TEM images of chloroplasts from VA-treated and non-treated plants at different timepoints immediately after dark night period. **c**, starch levels in different experimental variants immediately after dark period or after 4 hours of photosynthesis on light. **d**, acetate levels in control and VA-treated plants at different timepoints as determined by LC-MS analysis. Bars represent mean ±SD, with error bars showing variability among biological replicates. Data in (**a**, **c, d**) (*n* = 3). All statistical tests were carried out using a one-way ANOVA with post-hoc Tukey test with asterisk (APA guidelines) indicating significant differences (*P* < 0.05). Source data are provided as a Source Data file.

VA also redirects glycolytic pyruvate into acetate (Fig. 5a), an unusual flux shift first described by (Kim et al. 2017) that is central to jasmonate signaling and drought preparation. The VA-induced enzymes PDC1, ALDH2B4/7 and ADH1 are normally repressed by HDA6 (Kim et al. 2017). Accordingly, after 72 h VA-treated plants contained several-fold more acetate than controls (Fig. 5d), consistent with activation of HDA6-repressed acetate-mediated stress signaling.

### VA-induced metabolism reprogramming is accompanied by cell ultrastructural changes

Given the metabolic shifts and starch accumulation, the photosynthetic apparatus was examined ultrastructurally and functionally. GRANA neural-network analysis (Bukat et al. 2025) showed that, despite large starch granules, the thylakoid network remained intact and grana stacks were slightly larger (Fig. S3).

To test whether these changes affect photosynthesis, chlorophyll fluorescence was imaged at both timepoints (Fig. 6a-e). No differences appeared at 24 h; only after 3 days was there a small but significant drop in F_v_/F_m_ (0.78 vs 0.79 in controls). This drop in Fv/Fm was not accompanied by significant changes in the effective photochemical yield of PSII. This suggests that large starch granules lower maximum efficiency, while more compact grana compensate to keep effective photosynthesis unchanged. At 72 h, VA-treated plants also showed reduced non-photochemical quenching Y(NPQ) and increased non-regulated energy dissipation Y(NO), indicating reduced reliance on quenching-based protection against high light. These results are hard to interpret but may reflect the non-physiological starch accumulation stretching the chloroplasts, or that storage-filled cells deprioritize photosynthesis relative to building resistance.

**Fig. 6.**
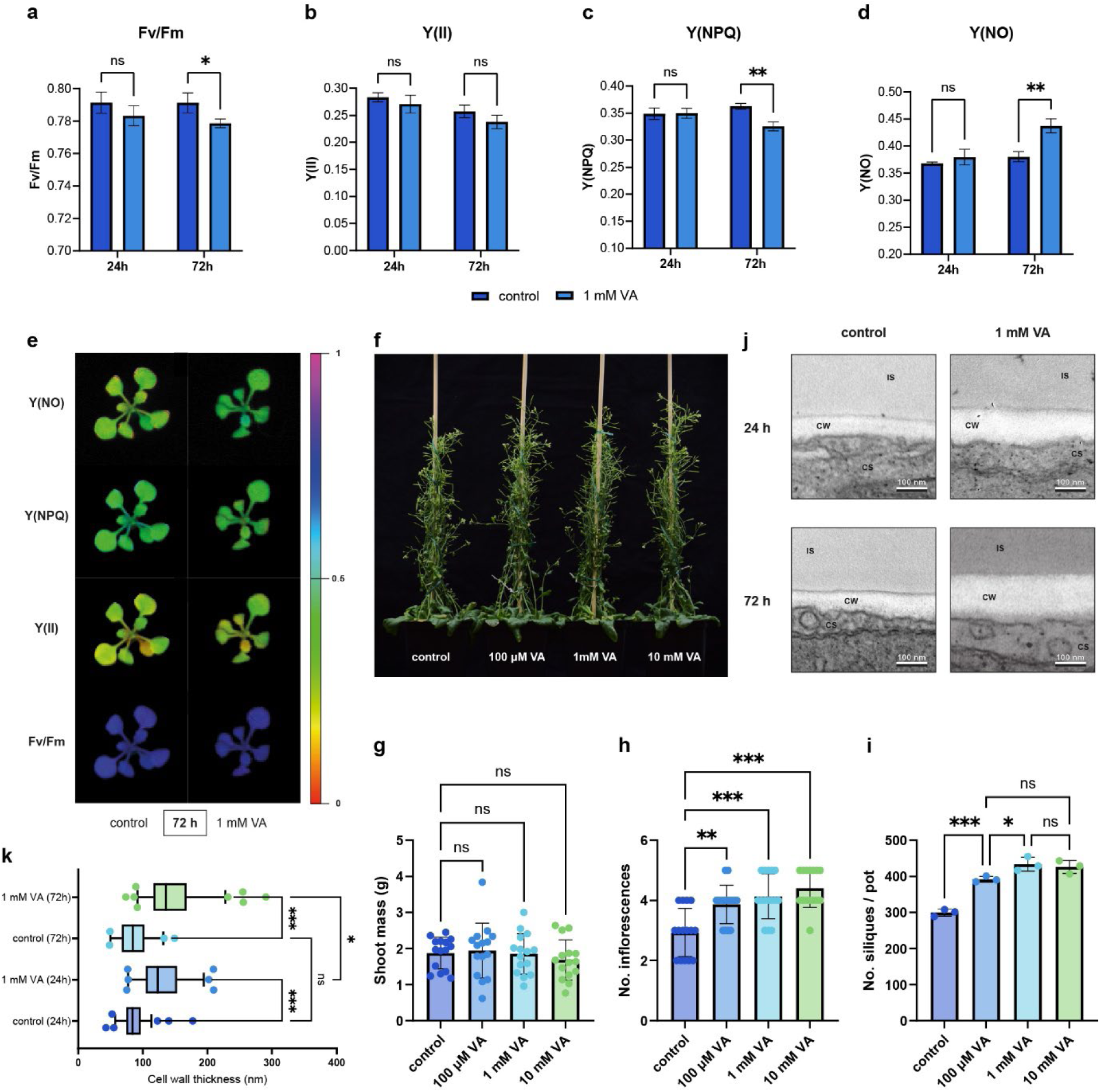
Metabolic reprogramming is accompanied by photosynthetic and generative development changes in VA-treated plants. **a**, **b**, **c**, **d**, photosynthetic parameters of control and 1 mM VA-treated plants at different timepoints showing maximal efficiency of PSII in the dark [F_v_/F_m_], effective photochemical yield of PSII [Y(II)], regulated non-photochemical yield of PSII [Y(NPQ)] and non-regulated non-photochemical yield of PSII [Y(NO)]. **e**, exemplary images of control and VA-treated plants at different timepoints showing the spatial distribution of photosynthetic parameters values. **f**, appearance of Arabidopsis plants in generative stage treated with VA at various concentrations. **g**, fresh mass of shoot of generative-stage plants from different variants. **h**, number of inflorescences per generative-stage plant from different variants. **i**, number of siliques per generative-stage plant from different variants. **j**, exemplary TEM images of the cell wall from control and VA-treated plants at different timepoints. **k**, thickness of cell wall in control and 1 mM VA-treated plants at different timepoints. Bars represent mean ±SD, with error bars showing variability among biological replicates. Data in (**a, b, c, d**) (*n* = 15), in (**g, h, i**) (*n* = 3, 5 plants per replicate), in (**k**) (control [24 h] *n* = 78, VA [24 h] *n* = 75, control [72 h] *n* = 42, VA [72 h] *n* = 85). All statistical tests were carried out using a one-way ANOVA with post-hoc Tukey test with asterisk (APA guidelines) indicating significant differences (*P* < 0.05). Source data are provided as a Source Data file.

Another carbohydrate sink, the cell wall, was also examined. It thickened markedly under VA, by ∼50% within 24 h and to ∼167% of control thickness by 3 days (Fig. 6j, k). This structural fortification possibly contributes to the multi-stress-resistant phenotype.

To test whether reduced photosynthesis and thicker cell walls affect productivity under non-stress conditions, greenhouse tests on mature plants were performed. Despite unchanged shoot mass, all VA-treated plants, at every concentration studied, produced more inflorescences per plant (Fig. 6f-h), indicating a shift toward generative over vegetative growth. Even 100 µM VA increased silique yield, and higher concentrations increased it further (Fig. 6i). Thus, despite broad defense priming, VA breaks the growth-defense trade-off.

## Discussion

This study shows that priming with valeric acid unlocks broad-spectrum resistance to abiotic and biotic stresses. Despite its simple structure, VA acts as a highly selective inhibitor of class I RPD3/HDA1 histone deacetylases and, as our data suggest, selectively targets HDA6 and HDA19 but not HDA9. This fine-tuning activates diverse defense mechanisms and leaves transcriptional memory for more efficient future responses. Additional extensive metabolic reprogramming that shifts flux toward gluconeogenesis and carbohydrate storage act as an energy reserve for stress. These findings address a central agricultural challenge: raising crop yield under worsening climate and more frequent extreme weather. The dominant approach, genetic engineering, is slow, expensive, and typically yields cultivars resistant to only one or a few stressors, while constitutive defense often imposes a growth penalty of stunted growth or lower yield (He, Webster, and He 2022). Biostimulants have emerged as a non-invasive alternative, but most are chemically complex or are plant-derived mixtures of unknown composition conferring a narrow resistance range (Arinaitwe, Yabwalo, and Hangamaisho 2025). VA is therefore a first-of-its-kind simple epigenetic biostimulant that induces multi-stress resistance without compromising growth, offering a way to address climate-driven crop losses from extreme weather and disease.

To our knowledge, this is one of the first reports of an HDA inhibitor conferring resistance to such a wide array of stressors; most HDAi studies focus on a single stress and miss the holistic effect. NaBT, the closest structural analog of VA, was shown in animals (Schölz et al. 2015) and here in plants to be class-I specific. Previously, only its benefit under salt and biotic stress was reported (Minoru Ueda et al. 2017; Y. Xu et al. 2022). VA differs from NaBT by only one - CH_3_ group yet inhibits HDA activity at lower concentrations (Fig. 1b). Moreover (Kim et al. 2017) showed that butyric acid does not raise *A.thaliana* resistance to drought, supporting VA’s uniqueness and intra-class I selectivity. The VA-primed phenotype resembles that of HDA6 or HDA19 knock-out mutants: VA-induced heat, drought, salt and biotic resistance maps onto the *hda19* mutant phenotype (Choi et al. 2012; M. Ueda et al. 2018; Minoru Ueda et al. 2017). HDA6 mutants likewise show elevated drought and biotic resistance, consistent with our findings (Kim et al. 2017; Y. Wang et al. 2017). Transcriptomic data indicate VA inhibits HDA6 and HDA19 but not HDA9, which is crucial for heat-signal transduction in Arabidopsis and whose inhibition by the non-specific pan-HDA inhibitor TSA increases heat sensitivity (Niu et al. 2022). The same TSA-induced heat sensitivity observed in this study (Fig. 2c) supports VA’s intra-class I selectivity. Not every aspect of the VA phenotype maps however onto HDA6/HDA19 inhibition. Freezing tolerance was never tested in *hda19* mutants nonetheless *hda6* mutants survive freezing far worse than wild type (To et al. 2011). Additionally, many developmental abnormalities present in *hda6* or *hda19* mutants were also not present in VA-primed plants.

HDA19 T-DNA mutants show retarded growth, drastically reduced seed yield, somatic embryogenesis and shoot-meristem abnormalities (Kotnik et al. 2025; Tanaka, Kikuchi, and Kamada 2008; Temman et al. 2023). HDA6 mutants show a similar phenotype, including delayed flowering (Wu et al. 2008; Yu, Chang, and Wu 2016). None of these developmental defects appeared in VA-primed plants, likely reflecting the recently described phenomenon of epigenetic moonlighting (Morgan and Shilatifard 2023). Many histone-writer/eraser enzymes, including HDA, act not only catalytically but also through the multivalent complexes they form (Chesnutt et al. 2025; Morgan and Shilatifard 2020). Because all data on plant HDA come from knock-out mutants lacking the full-length protein, it is unclear which traits depend on catalytic activity versus complex formation. VA impairs catalytic activity without removing the protein, and we propose that the remaining scaffolding function keeps developmental genes tightly controlled. Generating a double catalytic-dead HDA6/HDA19 mutant would evaluate whether the solely beneficial VA phenotype can be reproduced genetically.

The VA phenotype was also validated in monocots: drought and heat tests on maize showed that priming with VA drastically increased resistance to both. This fits the conservation of RPD3/HDA1 deacetylases across eukaryotes and especially plants (Alinsug, Yu, and Wu 2009). In rice, class I HDA also act in stress responses, and their inhibition or depletion enhances resistance to fungal infection or drought in *Oryza sativa* (X. Chen et al. 2025; Y. Xu et al. 2022). By targeting these evolutionarily ancestral mechanisms, VA is a versatile, high-potential biotechnological tool.

In the present study VA-primed phenotype was characterized on the molecular level. VA induced major stress-related transcriptomic reprogramming with supporting changes in primary energy metabolism, the 24 h response dominated by biotic-stress genes and abiotic ones less represented. Most of the biotic response was jasmonic acid (JA) signaling and its synthesis via fatty acid oxidation. This JA overstimulation follows from HDA6 inhibition, since HDA6 represses the acetate-mediated alternative JA signaling and biosynthesis pathway (Kim et al. 2017). HDA19 acts antagonistically to HDA6 on JA signaling (C. Zhou et al. 2005). As shown by (Vincent et al. 2022), the acetate-mediated JA pathway operates independently of standard signaling, so JA genes are upregulated in VA-primed plants despite HDA19 inhibition. For the SA-mediated response, also activated by VA, both HDA6 (Y. Wang et al. 2017) and HDA19 (Choi et al. 2012) repress SA-activated defense genes. It is consistent with present hypothesis and justifies why the VA-induced biotic response engages both JA and SA, which are normally mutually exclusive (Y. Wang et al. 2017). Together these explain the strong biotic response to VA. The less pronounced abiotic response, mainly ZAT transcription factors and oxidative- and heat-stress effectors, was nonetheless sufficient to confer the abiotic multi-stress resilience observed experimentally.

The time-course analyses show that VA treatment leaves transcriptional memory that primes future stress responses: almost half of the genes upregulated 24 h after heat shock had been primed by the initial VA treatment. Surprisingly, although the stress was heat shock, VA-primed plants strongly activated SA-mediated systemic acquired resistance (SAR), because most type-II memory genes (Oberkofler, Pratx, and Bäurle 2021) primed by VA were related to biotic response silenced by HDA6 and HDA19. This SAR activation, though unexpected, aided post-stress recovery, consistent with cross-tolerance (Kamran, Burdiak, and Karpiński 2025) and general defense activation. VA-primed type-I memory genes, by contrast, regulate long-term energy metabolism rather than stress response, showing that VA memory both speeds future responses and adapts metabolism to the energetic cost of defense. Given VA’s mechanism of targeted HDA inhibition, we propose that this memory depends on elevated histone acetylation at the loci of initially activated genes (Harris, Amtmann, and Ton 2023). Further ChIP-seq analyses are however needed to confirm this.

Metabolic reprogramming is central to the VA-resistant phenotype, with pyruvate/phosphoenolpyruvate metabolism and carbohydrate anabolism and storage at its core. Gluconeogenesis is driven by activation of the glyoxylate cycle, marked by high isocitrate lyase (ICL) expression (Cornah et al. 2004). Lately it has been shown that the glyoxylate cycle is activated in plants under stress and acts as a protective mechanism (Yuenyong et al. 2019), which aligns well with our findings. In VA-primed plants, using storage lipids and acetyl-CoA, it produces PEP and fuels gluconeogenesis, yielding surplus glucose imported into the chloroplast by the VA-induced GPT2 transporter, normally repressed by HDA19 (Shen et al. 2019), and stored as starch. This starch is degraded under stress, suggesting it serves as an energy reserve for costly defense responses, consistent with reports that stress-induced starch degradation benefits plants energetically and osmotically (Thalmann and Santelia 2017). The carbohydrate surplus may also supply the rapid cell-wall thickening seen under VA, which could explain the increased resistance to pathogens.

Despite the rewired metabolism, the photosynthetic apparatus stayed largely intact under normal conditions: although chloroplasts were packed with starch, the thylakoid network was preserved and grana became slightly more compact over time. Such grana architecture is typical of younger leaves (Gügel and Soll 2017) and increases photosynthetic efficiency (Gu et al. 2022), likely compensating for the obstruction caused by densely packed starch. Notably, the VA target HDA6 was recently found in chloroplasts, deacetylating many chloroplast proteins; its knock-out mutants, however, show a markedly different phenotype with a disrupted thylakoid network (H. Wang 2025), a discrepancy that again can be attributed to epigenetic moonlighting. The gradual drop in maximum quantum yield likely reflects chloroplasts reaching maximum starch-storage capacity.

Most importantly, under normal conditions VA increased yield without impairing whole-plant growth: VA-primed plants produced more inflorescences and siliques, showing that activated defenses do not penalize growth in soil. This may reflect VA-tuned water management combined with metabolic stimulation. It was unexpected, since HDA6 and HDA19 are crucial for flower development and their knock-out mutants show delayed flowering and developmental malformations (Gorham et al. 2018; Wu et al. 2008). This again shows that loss of catalytic activity does not simply mimic loss of the protein, underscoring epigenetic moonlighting in plant HDA.

Biotechnologically, these findings matter for future-proof crop protection based on native defenses, which are normally triggered by stress such as drought, heatwaves or infection. As climate change makes extreme events faster and more severe, a plant’s *ab initio* response might be too late to aid in adaptation, often leaving it vulnerable to severe damage or even death. Therefore priming agents like VA that allow for transient activation of stress response mechanisms are needed for modern agriculture. By targeting HDA6 and HDA19 that were shown to act as molecular switches of stress response, VA is able to prime the plants for stress. Moreover, by the fact of targeting epigenetic mechanisms, it is also able to leave stress memory that helps the plant to efficiently respond to future stress events. In the concept it is similar to the process of vaccination in animals (De Kesel et al. 2021) and is important for sustainable crop protection (Flors et al. 2024). VA is thus a first-of-its-kind targeted epigenetic biostimulant conferring both abiotic and biotic resistance, addressing a need recently postulated (Sako, Nguyen, and Seki 2021). Because of the metabolic reprogramming, VA priming does not impair growth and even raises yield. VA is also simple and natural, implying low cost and negligible ecotoxicity, and as a registered food additive it poses no consumer-safety risk on edible crops. Its rapid action also suits emergency use against forecast heatwaves or frosts, helping mitigate the climate-driven global crisis of crop losses from abiotic and biotic stress (Hultgren et al. 2025).

### Conclusions

This work sheds light on the previously largely unnoticed aspects and application potential of chemical inhibition of histone deacetylases in plants. It was shown that a molecule as simple as valeric acid can act as a selective inhibitor of class I HDA, putatively targeting only HDA6 and HDA19. This inhibition was demonstrated to unlock a broad range of abiotic and biotic stress resistance, not shown before for any other HDA inhibitor in plants. The transient stress response priming by VA was also able to break the defense to growth trade-off and increased crop yield in soil grown plants. Taken together with the stress memory induction by VA priming, the studied compound may be of high importance as a novel epigenetic biostimulant improving crop resistance and yield for future-proof agriculture. These findings also open new fields for discussion on the development of new highly selective HDA inhibitors for plant biotechnology and studies on non-catalytic functions of histone deacetylase in plants.

## Acknowledgements

This research was supported by the National Science Centre, Poland, under grant number 2019/35/B/NZ3/01362.

TEM studies were performed in the Laboratory of Electron Microscopy of the Nencki Institute, supported by the project financed by the Minister of Education and Science based on contract No 2022/WK/05 (Polish Euro-BioImaging Node “Advanced Light Microscopy Node Poland”).

## Author contributions

O.G conceived this study. O.G and M.K designed the experiments. O.G performed most of the experiments. M.G and O.G conducted biophysical analyses of VA binding to HDA. O.G, R.I-N, H.K and M.G performed the microarray experiments. O.G, P.Z and M.Krz designed and conducted biotic stress tests. O.G and R.M performed LC-MS analyses. Ł.K and OG conducted TEM ultrastructural analyses. O.G wrote the manuscript. M.K, Ł.K, K.W and O.G revised the manuscript. All authors discussed the results and contributed to the manuscript.

## Supplementary Figures

**Fig S1.**
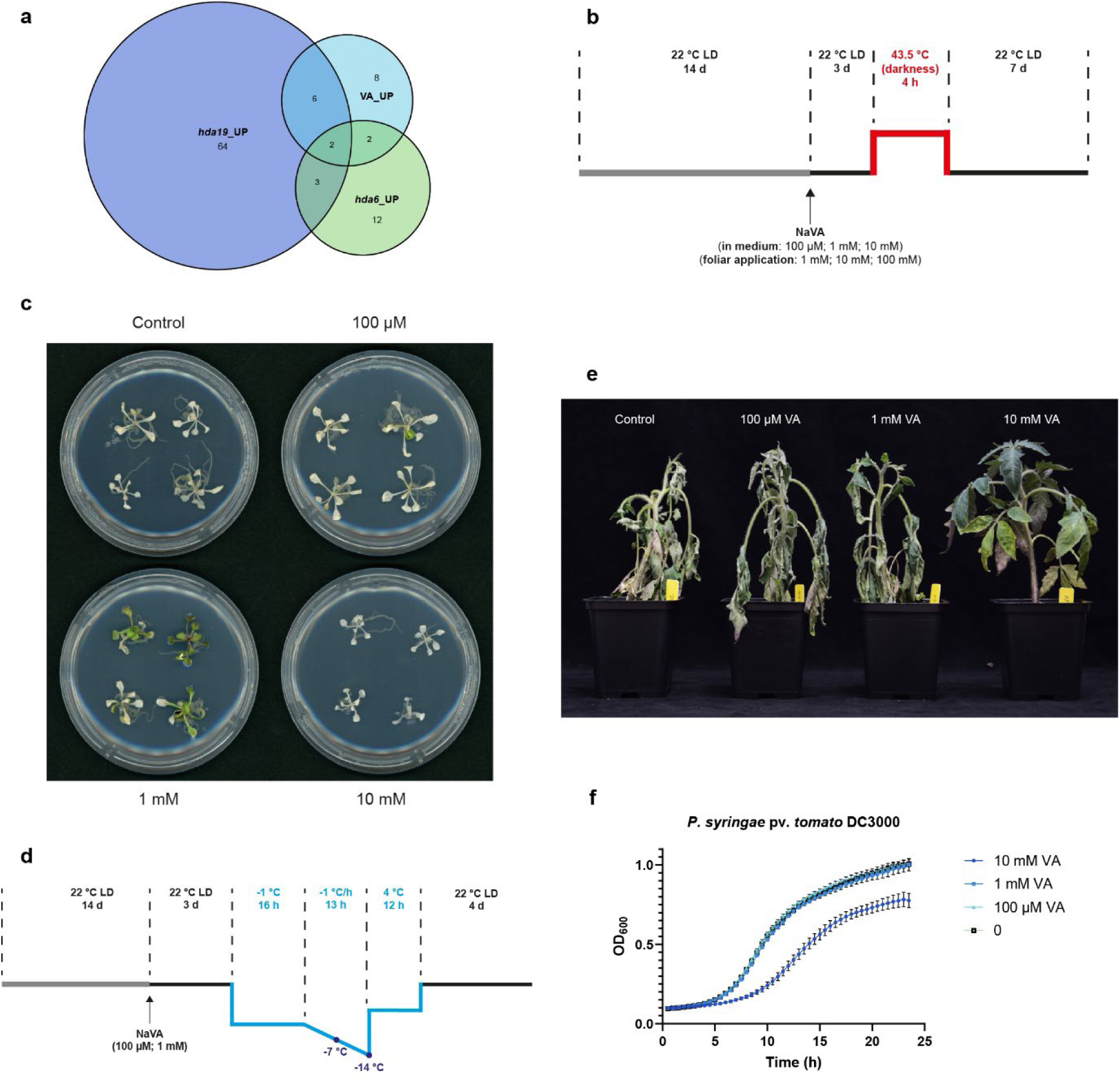
**a**, Venn diagram showing the overlap of upregulated transcription factors in VA-treated, *hda19* and *hda6* plants. **b**, schematic representation of heat stress experiments used in this study. **c**, appearance of Arabidopsis plants grown on MS media with various concentrations of VA 7 days after heat shock (4 h, 43.5°C). **d**, schematic representation of freezing stress experiments used in this study. **e**, VA-primed tomato (*Solanum lycopersicum*) plants after 7 days of drought period. **f**, *Pseudomonas syringae* pv. *tomato* DC3000 growth curves in the presence of different VA concentrations in the medium. Bars represent mean ±SD, with error bars showing variability among biological replicates. Data in (**a**, **f**) (*n* = 3), in (**e**) (*n* = 4). All statistical tests were carried out using a one-way ANOVA with post-hoc Tukey test with asterisk (APA guidelines) indicating significant differences (*P* < 0.05). Source data are provided as a Source Data file.

**Fig. S2.**
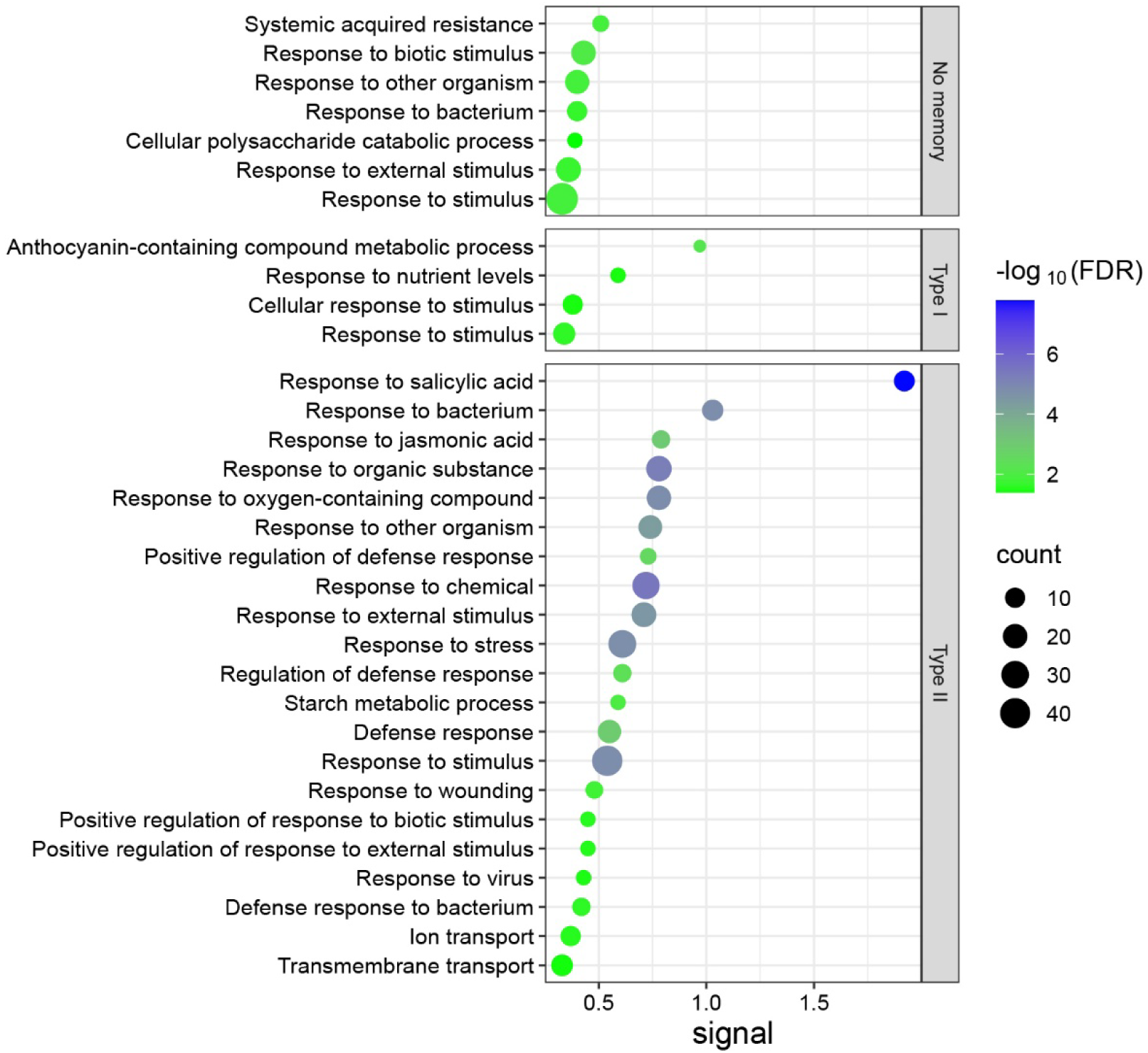
GO enrichment analysis of upregulated genes showing transcriptional memory in VA-primed plants 24 h after heat shock.

**Fig. S3.**
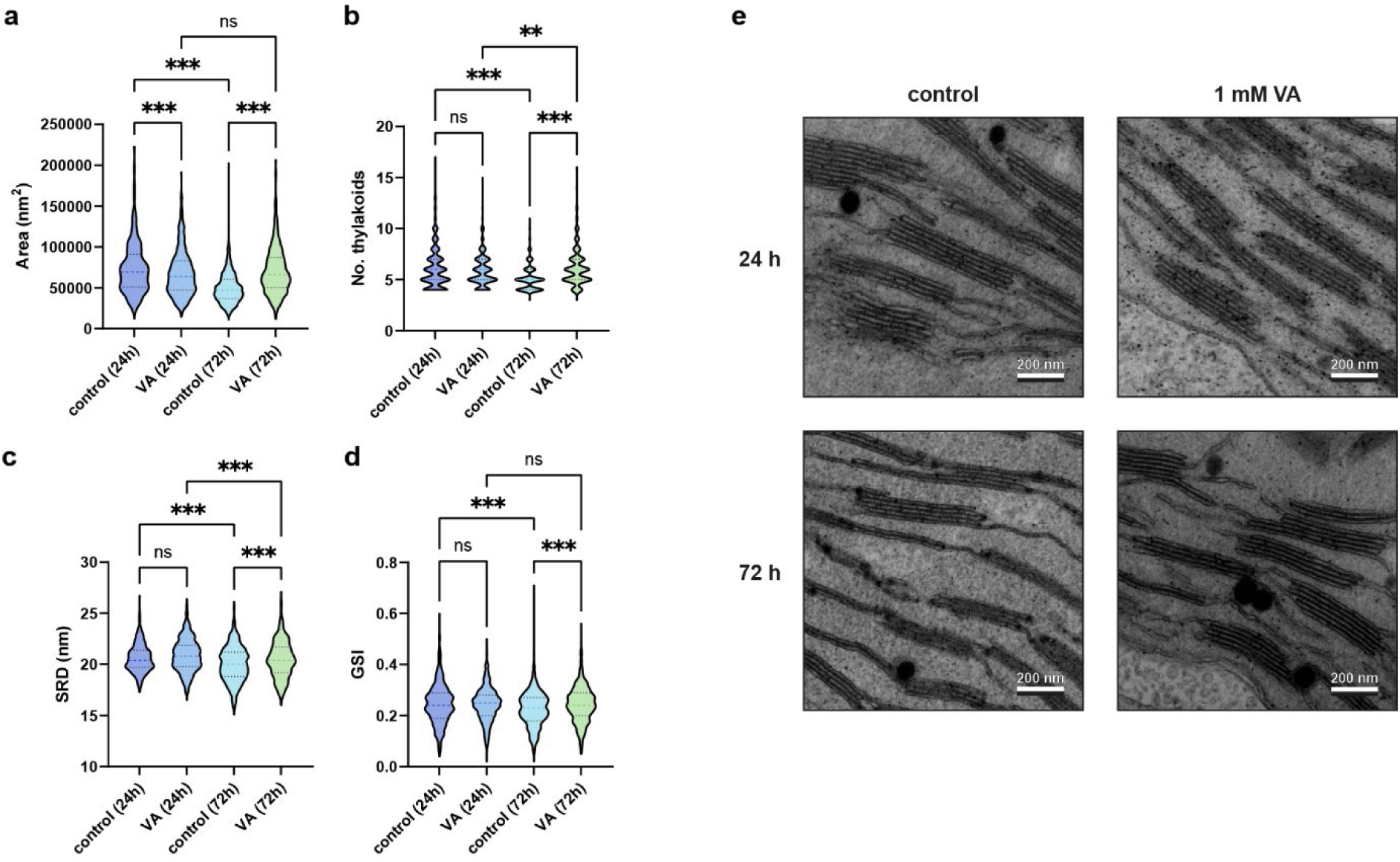
**a-d**, results of grana morphology analysis performed in GRANA showing thylakoid area, number of thylakoids per granum, thylakoid stacking repeat distance [SRD] and grana cross-sectional irregularity [GSI] in control and 1 mM VA treated plants. **e**, exemplary TEM images of thylakoid networks from different experimental variants. Bars represent mean ±SD, with error bars showing variability among biological replicates. Data in (**a, b, c, d**) (area: control [24 h] *n* = 765, VA [24 h] *n* = 1250, control [72 h] *n* = 1952, VA [72 h] *n* = 884; no. thylakoids/SRD: control [24 h] *n* = 387, VA [24 h] *n* = 533, control [72 h] *n* = 932, VA [72 h] *n* = 469; GSI: control [24 h] *n* = 619, VA [24 h] *n* = 950, control [72 h] *n* = 1607, VA [72 h] *n* = 619;). All statistical tests were carried out using a one-way ANOVA with post-hoc Tukey test with asterisk (APA guidelines) indicating significant differences (*P* < 0.05). Source data are provided as a Source Data file.

